# Pavlov’s pea plants? Not so fast. An attempted replication of Gagliano *et al*. (December 2016)

**DOI:** 10.1101/2020.04.05.026823

**Authors:** Kasey Markel

## Abstract

Gagliano *et al*. (Learning by association in plants, 2016) reported associative learning in pea plants. Associative learning has long been considered a behavior performed only by animals, making this claim particularly newsworthy and interesting. In the experiment, plants were trained in Y-shaped mazes for three days with fans and lights attached at the top of the maze. Training consisted of wind consistently preceding light from either the same or the opposite arm of the maze. When plant growth forced a decision between the two arms of the maze, fans alone were able to influence growth direction, whereas the growth direction of untrained plants was not affected by fans. Importantly, some plants were trained to grow towards the fan and others to grow away, demonstrating the flexibility of associative learning. However, a replication of their protocol failed to demonstrate the same result, calling for further verification and study before mainstream acceptance of this paradigm-shifting phenomenon. This replication attempt used a larger sample size and fully blinded analysis.

## Introduction

The temporal and spatial heterogeneity of any given environment poses a major challenge to all life on Earth. Accordingly, a common characteristic of living organisms is the capacity to respond to environmental cues. Many of these responses involve nuanced decisions integrating multiple cues or evaluating cue intensity, such as chemotaxis in bacteria and other microbes (Wadhams and Armitage, 2004). Even ‘simple’ organisms make decisions based on the integration of multiple cues or the intensity of particular cues. For example, unicellular *Dictyostelium discoideum* amoebae aggregate into multicellular slugs in the absence of food, and travel at precise, genetically determined angles in the direction of a light source (Annesley and Fisher, 2009).

Many complex environmental responses have recently been discovered in plants, including luring in animal predators to attack herbivores in response to herbivory (De Moraes *et al*., 1998), production of toxins in response to distress signals from kin (Karban *et al*., 2013), and dose-dependent responses to insufficient water (Pandey *et al*., 2016) or excess salt (Julkowska and Testerink, 2015) in the soil. These responses demonstrate an impressive capacity to detect a variety of environmental signals and respond accordingly. These instinctive behaviors have been sculpted over evolutionary time, and can be contrasted with learned behaviors that are developed within the lifetime of an individual as a result of their particular interactions with the environment. Individuals and groups of organisms leverage learning to exploit patterns in their environment over short timescales, and the capacity to do so frequently confers a selective advantage (Morand-Ferron, 2017). Learning is useful in any environment that contains exploitable regularities and patterns and for which the instincts of the organism have not evolved to perfectly exploit those patterns. Learning is ubiquitous among animals, from zebrafish (Sison and Gerlai, 2010) and insects (Prokopy *et al*., 1982) to nematodes (Wen *et al*., 1997), and recent work in *Caenorhabditis elegans* is beginning to unravel its molecular and genetic mechanisms (Gyurkó *et al*., 2015).

Over the last decade, there has been substantial debate in neurobiology and philosophy of mind regarding what kinds of organisms possess the capacity to learn (Gagliano *et al*., 2016, 2018; Thellier, 2017; Gagliano, 2017), along with thornier issues like cognition (Garzón, 2007; Gross, 2016; Adams, 2018; Segundo-Ortin and Calvo, 2019) and intelligence (Marder, 2013; Trewavas, 2017). Some of these disputes center around particular terminology and language, but some basic empirical questions remain contentious, including the simple question: can plants learn? A series of experiments on associative conditioning have been carried out in plants since the 1960s, notably the efforts of Holmes and Gruenberg (Holmes and Gruenberg, 1965), Haney (Haney, 1969), Levy *et al*. (Levy *et al*., 1970), and Armus (Armus, 1970). These studies generally failed to observe conditioning in plants, and those that concluded in favor of conditioning lacked sufficient controls to rule out other explanations, as reviewed by Adelman (Adelman, 2018). Gagliano *et al*. (Gagliano *et al*., 2016) provide the most convincing report of conditioning in plants to date, but there are no published reports of replication, from the original lab or others.

Associative learning is the phenomenon whereby an individual organism associates two stimuli, and thereafter uses one as an indicator for the other. In Ivan Pavlov’s classic work, dogs were trained to associate the ringing of a bell with the imminent arrival of food, and demonstrated the appropriate physiological response of salivation (Pavlov, 1927). Associative learning experiments attempt to pair an unconditioned stimulus for which there is an observable response to a conditioned stimulus which is originally neutral. Learning is demonstrated when presentation of the conditioned stimulus can elicit responses normally associated with the unconditioned stimulus. In December 2016, Gagliano *et al*. published a report claiming plants have the capacity to learn adaptive behaviors during individual lifetimes using an experimental setup analogous to Pavlov’s famous experiment (Gagliano *et al*., 2016).

## Methods

*Pisum sativum* cv Green Arrow seeds were germinated in the dark following the protocol of Gagliano *et al*. After germination, seedlings were planted in round pots in a growth chamber (8:16 light:dark cycle, 20 °C, 85% humidity) until emergence from soil. An excess number of plants were planted to allow for selection of plants at a relatively homogenous growth stage and central positioning within the pots. The Y-maze apparatus was then attached, and only thereafter were plants randomly (using a random number generator) divided into experimental conditions to prevent bias. Each experimental condition was then split into two opposite versions starting on the right or the left but displaying the same pattern. These symmetrically paired conditions were grouped for analysis, following Gagliano *et al*. Once the Y maze was attached to each pot, plants were moved into a dark chamber with the same humidity, and training commenced. Once within this chamber, all plant manipulation took place with a dark external room under a dim (∼0.8 lux measured at light source, <0.2 lux measured at seedlings) red LED light. Training consisted of 3 condition-specific combinations of blue LED lights and 35-mm fan-generated wind presented in the two arms of the Y-maze per day, as diagrammed in Figure 1. After three days of training, all plants which had already grown past the ‘decision point’ in the maze were disqualified, and the testing day began, in which plants were presented only with wind, no light. Plants were then removed from chambers, the lights and rans removed leaving only the Y-maze structural core, randomly arranged along the bench, and scored by observers with no other connection to the experiment with the experimenter in a separate room. Scoring criteria were simple and objective: growth >5 mm above the decision point in only one maze arm. Failure to grow in either maze arm or growth into both maze arms was scored as disqualification. Additional details including all suppliers, part numbers, and detailed description of apparatus assembly are available in supplemental methods and materials.

**Figure 1:**
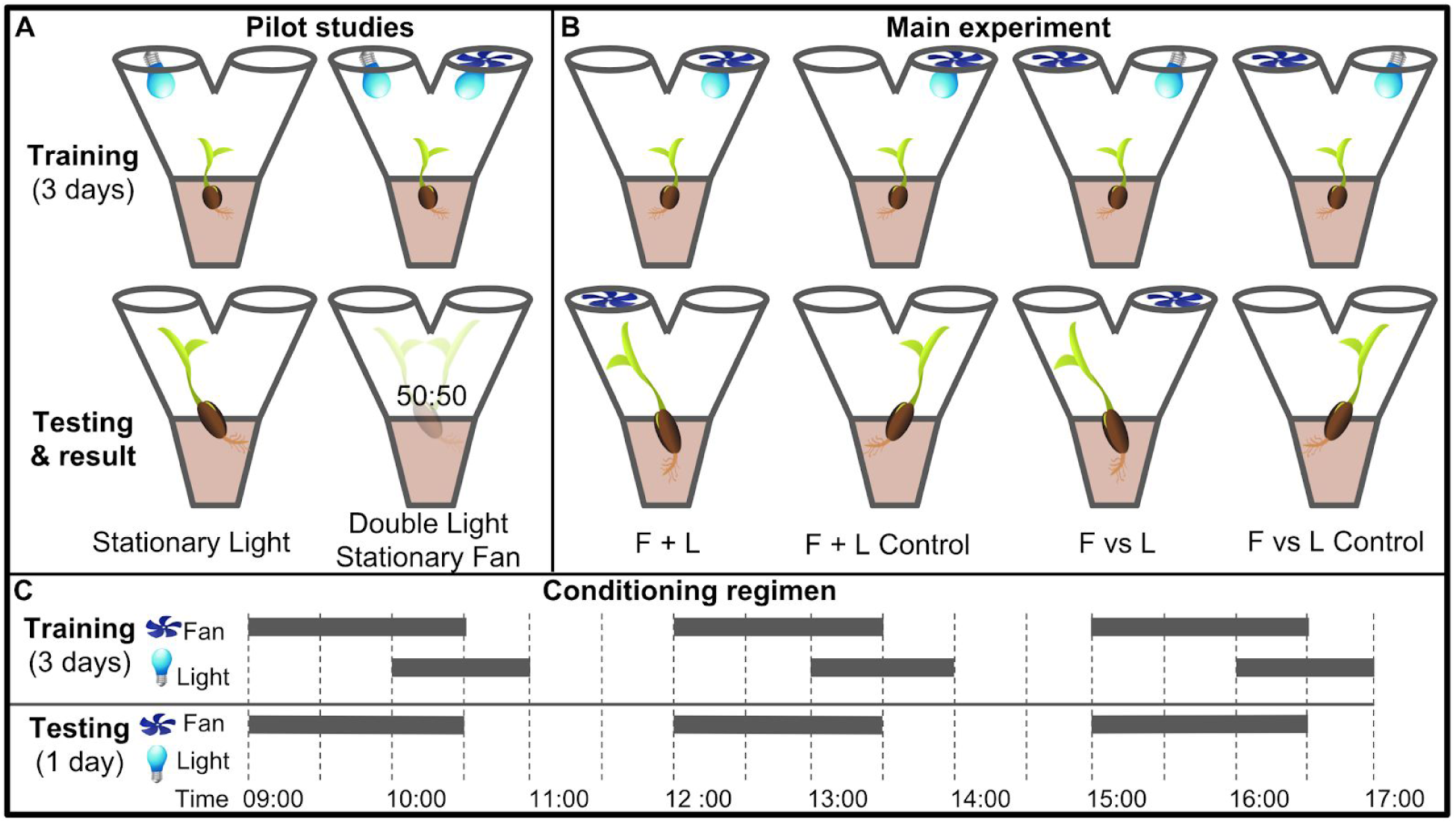
Visual summary of the two Gagliano *et al*. pilot studies and four experimental conditions. A: Pilot studies used. Top row shows maze fan and light configuration for training period, bottom row shows configuration during testing day and the maze arm into which plants grew. B: Main experimental conditions for “Experiment 1”. Top row shows maze fan and light configuration for training period, bottom row shows configuration during testing day and the maze arm into which most plants grew. C: Conditioning regimen. Horizontal bars indicate time periods for which fans and lights were active, the x-axis indicates time of day.

## Results

### Summary of original study

Gagliano *et al*. adapted Pavlov’s paradigm for pea plants (*Pisum sativum)* using light as the unconditioned stimulus and wind as the neutral stimulus, which through training became the conditioned stimulus (Gagliano *et al*., 2016). Plants were grown inside a dark controlled growth chamber with individual PVC Y-mazes with blue LEDs and fans attached on each arm, and plant learning was inferred through the maze arm selection of the plants. This experimental setup is particularly suited to pea plants, which are topped during their early stages by a single tendril, which makes maze arm selection quite clear. See supplemental methods for growth conditions, germination protocol, and maze dimensions, which are unchanged in this study. Supplemental figure 1 details the construction of the Y maze and the associated lights and fans required for the experiment.

For the main experiment, seedlings were randomly assigned to two groups: F vs L presented light and fan on opposite maze arms, F + L presented light and fan on the same arm. The plants were trained for three days with three exposures per day with wind proceeding light by 60 minutes, overlapping for 30 minutes, then light only for an additional 30 minutes. These exposures were moved from one arm to the other according to the pattern: day 1, left (L)/right (R)/L; day 2, L/R/R; day 3 R/L/L, tested on R, or the inverse (plants were randomly assigned to opposite patterns and grouped for analysis). These three days of training were timed with the plants’ growth such that on the fourth day the plants would grow into one arm or the other of the maze. Lights were disabled for this day, and the plants were subdivided into control and experimental groups.

Control plants were left undisturbed and grew towards the arm where light was last presented on day 3, whereas experimental plants had fans placed so as to suggest light would be presented from the opposite arm. Gagliano *et al*. report that approximately two thirds of experimental plants showed behavior influenced by the fan, thus demonstrating associative learning. The setup and results of the two pilots and four main experimental conditions are shown in Figure 1.

Gagliano *et al*. also performed two pilots: Stationary Light and Double Light Stationary Fan.

In stationary light, a LED was fixed on one arm and turned on for three one-hour periods per day for three training days, then turned off for the testing day. All plants grew into the arm with the LED, confirming positive phototropism. In Double Light Stationary Fan, lights were affixed to both arms and a blowing fan was fixed on one arm. Plants grew at random, unaffected by the fan. Taken together, these pilots showed light is a suitable unconditioned stimulus and wind is a suitable candidate conditioned stimulus, at least when both stimuli are stationary.

### Methodological changes

The Gagliano *et al*. experiment was clever and innovative, and reports a finding of outstanding interest. Findings this unexpected have the potential to open up a new field of study, the establishment of which requires independent verification and experimental rigor (Kuhn, 2012). This section details departures from the original study design, which consist of controls, sample size, and blinding.

The two Gagliano *et al*. pilot studies involved stationary stimuli, while the main experiments and interesting findings involve moving stimuli. Therefore, we chose a different set of pilot controls, designated Moving Light, Moving Light Fan Opposite, and Moving Light Fan Adjacent. The training period for all three of these consisted of a light moving between maze arms in the same pattern as the lights in the four main conditions. On testing day, Moving Light was given no stimulus, Moving Light Fan Opposite had a fan placed opposite the last light exposure, and Moving Light Fan Adjacent had a fan placed on the same maze arm as the last light exposure. Moving Light is intended to confirm that the training regimen consistently results in growth towards the last presentation of light in the absence of wind, the other two are to check for systematic growth towards or away from the fan as well as any “fan-induced amnesia,” wherein the fan interferes with the plant’s growth towards the last presentation of light, an alternative explanation that might account for the Gagliano *et al*. results. Importantly, the four main conditions of the study, F + L, F + L Control, F vs L, and F vs L control, were performed identically to the original experiment.

Sample size in the Gagliano *et al*. study was sufficiently large to generate strong evidence, with several p values under 0.005. The present study increased sample size significantly to allow detection of a weaker effect and estimate the effect size more precisely. The blinding protocol is also more complete. In the original study the two experimenters also performed the scoring: the experimenter who set the experiment up recorded while the other scored the plants. In this replication attempt, scoring began with the experimenter removing all plants from the growth chamber, removing the fans and lights, and placing plants still within Y-mazes in a random order along the bench. Thereafter, an independent observer who had no other involvement in the experiment scored the plants and recorded scores while the experimenter was in a separate room. The scoring protocol was precisely defined as follows: if any part of the plants has grown more than 5 mm into a maze arm the plant was scored as choosing that arm, unless there was any growth into the other arm. If any growth was present in both arms or if <5 mm growth was present in either arm, plants were scored as choosing neither and not counted.

### Comparison of results

The controls Moving Light Fan Opposite and Moving Light Fan Adjacent were used to test for an inherent attraction or repulsion to the fan stimulus, and found no significant effect on growth direction (p=0.1713, Fisher’s exact test (Fisher, 1935), two-tailed). This suggests fans are an acceptable candidate conditioned stimulus. However, while Gagliano *et al*. demonstrated plants always grow towards a stationary light, we chose to ask whether plants always grow towards the last presentation of a light moving in the same regimen as the experiment, the Moving Light condition. Their growth did not differ significantly from a random 1:1 expectation at this sample size (p=0.5885, Fisher’s exact test). The results of the pilot studies in this study and the Gagliano *et al*. study are compared in Figure 2.

**Figure 2:**
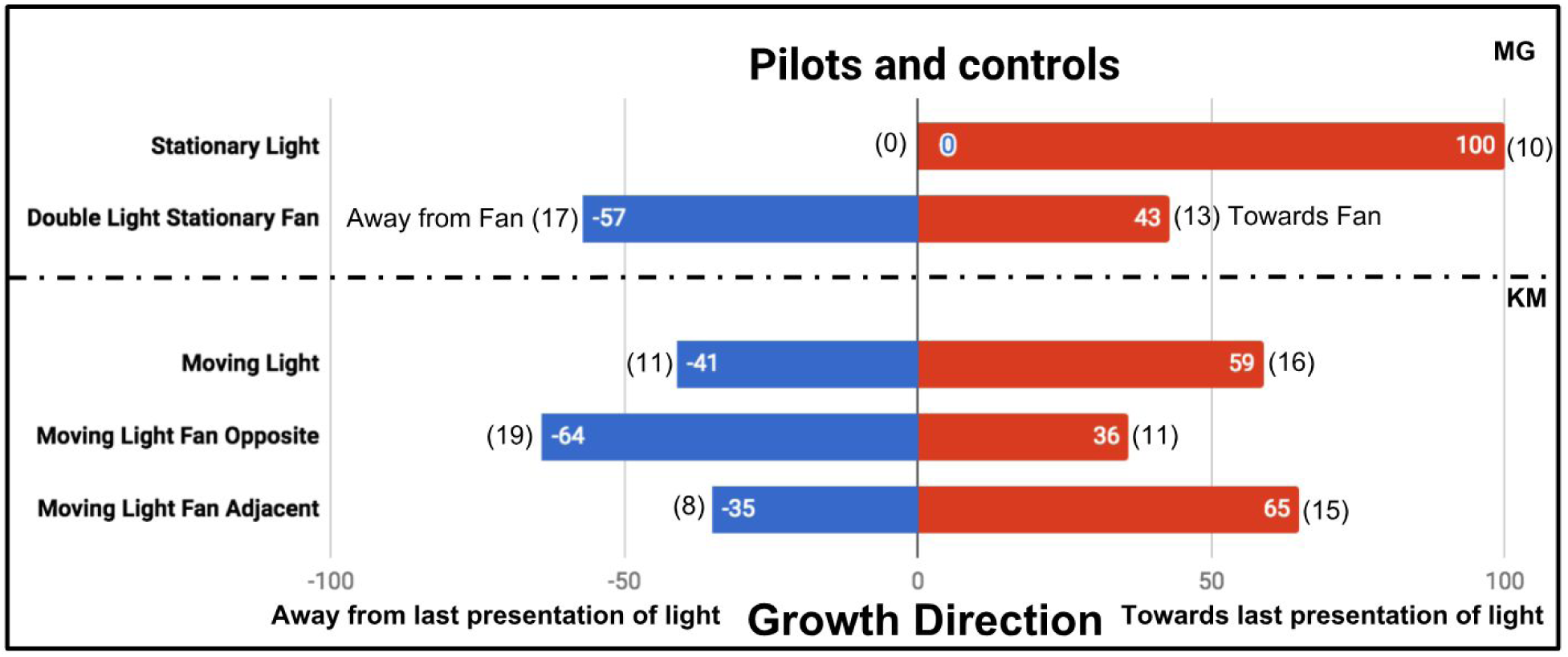
Results of plant growth in control conditions from the present study (KM) and the Gagliano *et al*. 2016 experiment (MG). Numbers in bars indicate percentage, numbers in parentheses indicate raw data, number of plants growing into each maze arm.

In the four main conditions the difference in results is fundamentally the same. F + L Control and F vs L Control plants did not always grow towards the last presentation of light, though interestingly both, like the Moving Light control, had a slight tendency in that direction. This tendency was statistically significant only in F + L control (also the condition with the largest sample size, n=61, p=0.0243). In other words, there is a tendency to grow towards the last presentation of light, even if the presentation of light was very brief (1 hour) and proceeded by presentations of light from the opposite direction. However, we found this effect to be much weaker than the perfectly phototropic response Gagliano *et al*. report, despite using very similar lighting.

In the F + L Control and F vs L Control, we also found a slight tendency to grow towards the last presentation of light but far weaker than the perfect directionality reported by Gagliano *et al*. The noise in the control conditions makes any associative learning more difficult to detect. While Gagliano *et al*. report a significant difference between F + L and F + L Control (p=0.0027, n = 13 + 10, Fisher’s exact test, two-tailed), we found no significant difference (p=0.335, n=61 + 60). Similarly, Gagliano *et al*. report a significant difference between F vs L and F vs L Control (p=0.0017, n= 13 + 9), whereas we found no significant difference (p=0.387, n=42 + 40). The results of the main four conditions are shown in Figure 3.

**Figure 3:**
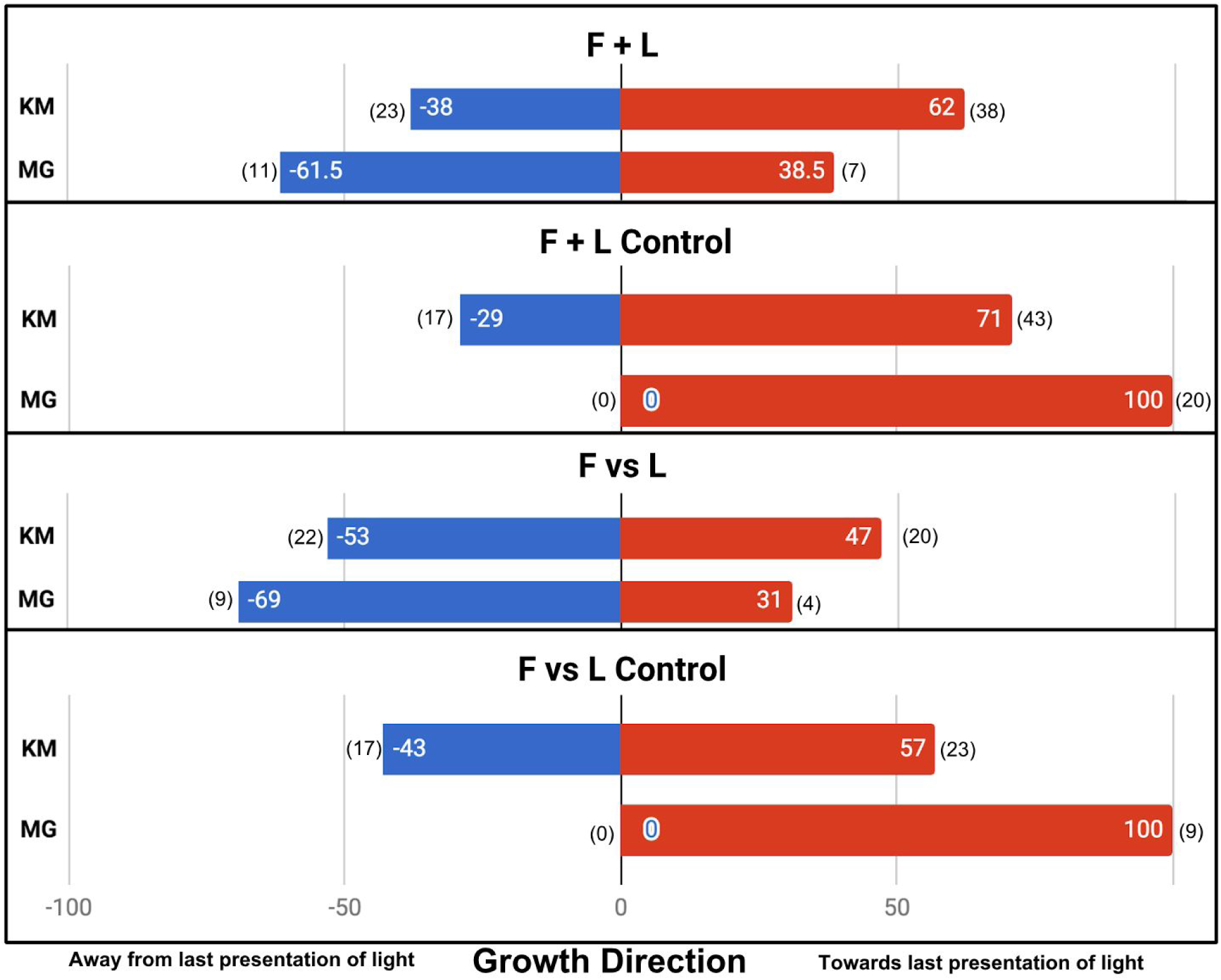
Results of plant growth in the 4 main conditions from this experiment (bars labeled KM) and the Gagliano *et al*. 2016 paper (bars labeled MG). Numbers in parentheses indicate sample size. Data are pooled from the Gagliano *et al*. “Experiment 1” and the Light group of their “Experiment 2”.

## Discussion

Gagliano *et al*. reported that in the absence of fan stimuli, pea plants always grow towards the last one-hour presentation of light, whereas we find their growth to be only slightly biased towards the last presentation of light. It is possible that this difference is due to the use of different cultivars of *Pisum sativum -* Gagliano *et al*. used *cv* Massey gem, whereas this study used *cv* Green Arrow because Massey Gem was not available. These cultivars are closely related, are both full sun varieties, and have a similar growth habit. There were several other minor differences in materials used, but importantly growth conditions, maze construction, training regimen, and fan and light intensity were identical to the conditions reported by Gagliano *et al*. Because the capacity for associative learning is a complex trait, it is expected to not have evolved in the few decades since these cultivars shared a common ancestor, and seems equally unlikely to have been lost from one cultivar but not another in the absence of specific selection. However, a stronger phototropic response in Massey Gem cannot be excluded, calling for further replication efforts.

We hope to see these performed using larger sample sizes and fully blinded scoring, and ideally with input and assistance from the original authors. Input from the original authors would allow for a more accurate replication, such as using the same pea plant supplier, LEDs, and fans, which may be critical to reproduce the phenomenon. If these efforts reproduce the phenomenon, we predict this experimental system will become widely used in experiments on plant learning. One shortcoming in the experimental system is that variation in plant growth rates results in substantial attrition. Plants were examined the morning of testing day to determine if they had already grown into one arm of the maze, and if they had they were excluded from the analysis. Furthermore, after testing day some plants had still not grown into either arm, and were therefore also excluded from the analysis. While care was taken to grow plants as uniformly as possible, approximately 40% of the plants were disqualified for growing too fast or too slow, as shown in Table 1.

**Table 1:**
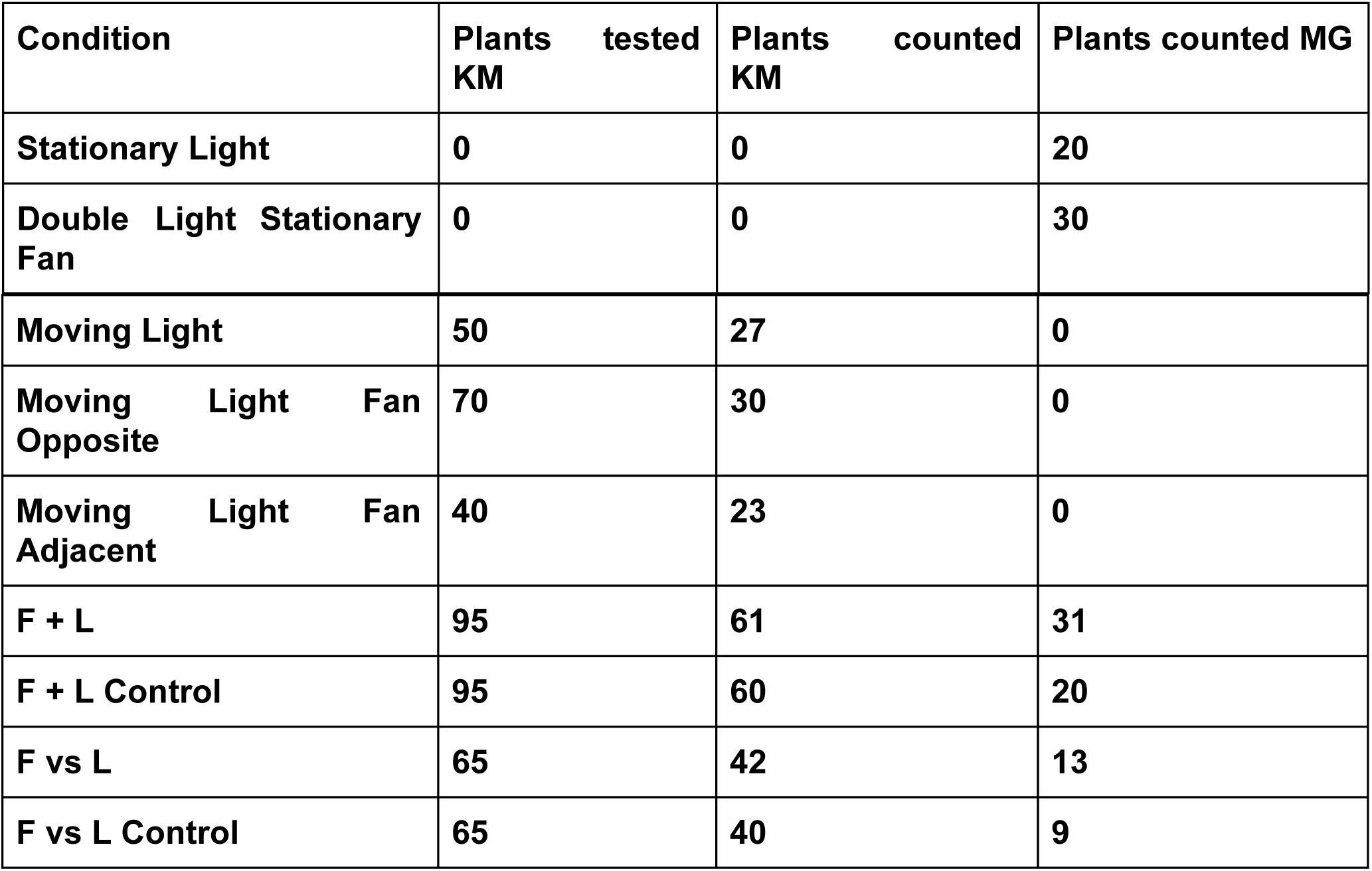
Sample size summary table. This experiment (Columns labeled KM) suffered significant attrition due to variance in plant growth rate. Sample sizes from the Gagliano *et al*. 2016 paper (MG) are shown for comparison, grouping results from their “Experiment 1” and the light condition of “Experiment 2”. Raw data available in Supplemental Datasheet.

The December 2016 report of associative learning in plants has garnered substantial attention in the press (*Smarty Plants* | Radiolab | WNYC Studios; Morris, 2018; robby-berman, 2018), and the reported phenomenon is extremely interesting. Gagliano *et al*. conclude this phenomenon will force us to reconsider the nature of learning (Gagliano, 2017) and we agree, contingent on the phenomenon being reproducible. Associative learning in the absence of traditional neurons must have a fascinating molecular mechanism and would open a new field of research. Unfortunately, no reports of successful replication have been published since the initial paper. Furthermore, this attempt at replication casts some doubt on the underlying premise used for scoring plant response, namely that plants will always grow towards the most recent one-hour light exposure after repeated exposures on different arms of a Y maze. At the least, it suggests that the conditions required for the experimental setup to function properly may be more precise than the stated parameters in the original paper. Additional work from the original authors and replications in independent labs are needed if this fascinating phenomenon is to move into the scientific mainstream.

## Supplemental information

**Supplemental materials and methods:** Details of plant growth conditions and apparatus setup.

**Figure S1:** Y maze assembly process and the final growth chamber with 120 Y mazes inside.

**Supplemental data:** Raw data for all experiments performed and statistical summary.

### Data availability

All data generated or analyzed during this study are included in this article and its Supplementary Information files.

## Acknowledgements

I would like to thank Andrew Zink and Robyn Crook for their ideas regarding study design, Thomas Varley and Rebecca Dumanski for critical reading of the manuscript, Karen Markel for scoring plants, Anireddy Reddy for access to controlled growth chambers, Kyaw Tha Paw U for assistance with wind speed measurements, and Monica Gagliano for the original report and innovative experimental design.

## Author Contributions Statement

K.M. designed and performed the experiments, analyzed results, and wrote the manuscript.

