## Supplemental Materials and Methods for "Pavlov’s pea plants? Not so fast. An attempted replication of Gagliano *et al*. (December 2016)"

**Supplemental methods**

To accompany the manuscript

**Pavlov's plants? Not so fast.**

An attempted replication of Gagliano *et al.* (December 2016)

Kasey Markel

### Germination conditions

Peas (*Pisum sativum* cv Green Arrow, Botanical Interests, USA) were germinated hydroponically in round containers kept in the dark. Seeds were first soaked in water for 24 hours, then wrapped by wet paper towel surrounded by aluminum foil in vertical rolls. These rolls were placed in water and incubated in the dark at 20 °C, changing the water daily. Roughly three times as many seeds were germinated as plants desired, and after 5 days of water incubation seeds were inspected for germination using a dim red LED headlamp (~0.8 lux measured at light source, <0.2 lux measured at seedlings). Seeds with radicle >5 mm were considered to be germinated, ungerminated seeds were discarded. Among the germinated seeds, those with particularly long or short radicles were discarded to minimize variance in growth stage. Roughly twice times as many germinated seedlings were kept as plants desired. Each seedling was then planted in the center of round pots (5 cm diameter at top, 6 cm deep, 4 cm diameter at bottom, with a single drain hole of ~2 mm diameter in the center) in Hoffman seed starter potting and planting mix (Good Earth Inc, NY, USA), at a depth of 15 mm. Soil was first watered to saturation, seeds were planted, and plants watered again to saturation. Soil was allowed to drain for ~30 minutes, then germinated seedlings were moved into a controlled growth chamber (PGR15 Growth Chamber, Conviron) until emergence from soil.

### Growth conditions

Chamber was maintained at 20 °C, 85% humidity, with blue and red LEDs balanced for an approximation of white light. Light was delivered at 50 μmol $m^{-2}$ $s^{-1}$ at soil surface with 8:16 hour light:dark cycle with light phase beginning at 09:00 (identical to Gagliano *et al.*). After 3-4 days, most seedlings had emerged from soil. Seedlings at as similar of a growth stage as possible were selected, watered to saturation and allowed to drain, and the Y maze attached. To correct for some plants being minorly off-center, all plants were rotated to maximize left-right symmetry before the attachment of the maze (importantly, the relevant growth directions had not been assigned at this stage, so there is no possible bias in the attachment of mazes, which were as centered as possible). Once all plants had mazes attached to their pots, they were assigned to groups using a random number generator, then grouped in rows and placed into the experimental growth chamber, where the fan and LEDs were attached on top of the maze arms.

**Apparatus details**

The experimental apparatus consisted of blue LEDs with light intensity ~14 μmol $m^{-2}$ $s^{-1}$ (measured at soil surface) in the 430–505 nm wavelength range (CO RODE part#CR150514E156, Dr. Gagliano did not specify a particular LED model and did not respond to requests for model/part numbers). Fans were 35 mm 10,000 RPM computer cooling fans (Gdstime XH2.54-2Pin 3510S 35x10mm brushless fan). The fans generated a semi-turbulent airflow of 0.6-0.8 m $s^{-1}$ at all points in the Y maze, including soil surface, the branching point of the maze, and on the fan arm and the opposite arm. Flow was approximately downward in the fan arm (parallel to the arm of the maze) and upward in the opposite arm, and was measured with TPI 575 anemometer hotwire probe and vane, which each gave similar results. These systems were soldered to two sets of 12V DC wires which extend throughout the growth chamber, and are powered by a digital automated timer system. 12V DC power was generated from 120 V 60 Hz AC power by a Winkeyes power supply (ASIN# B018G3ABWY).

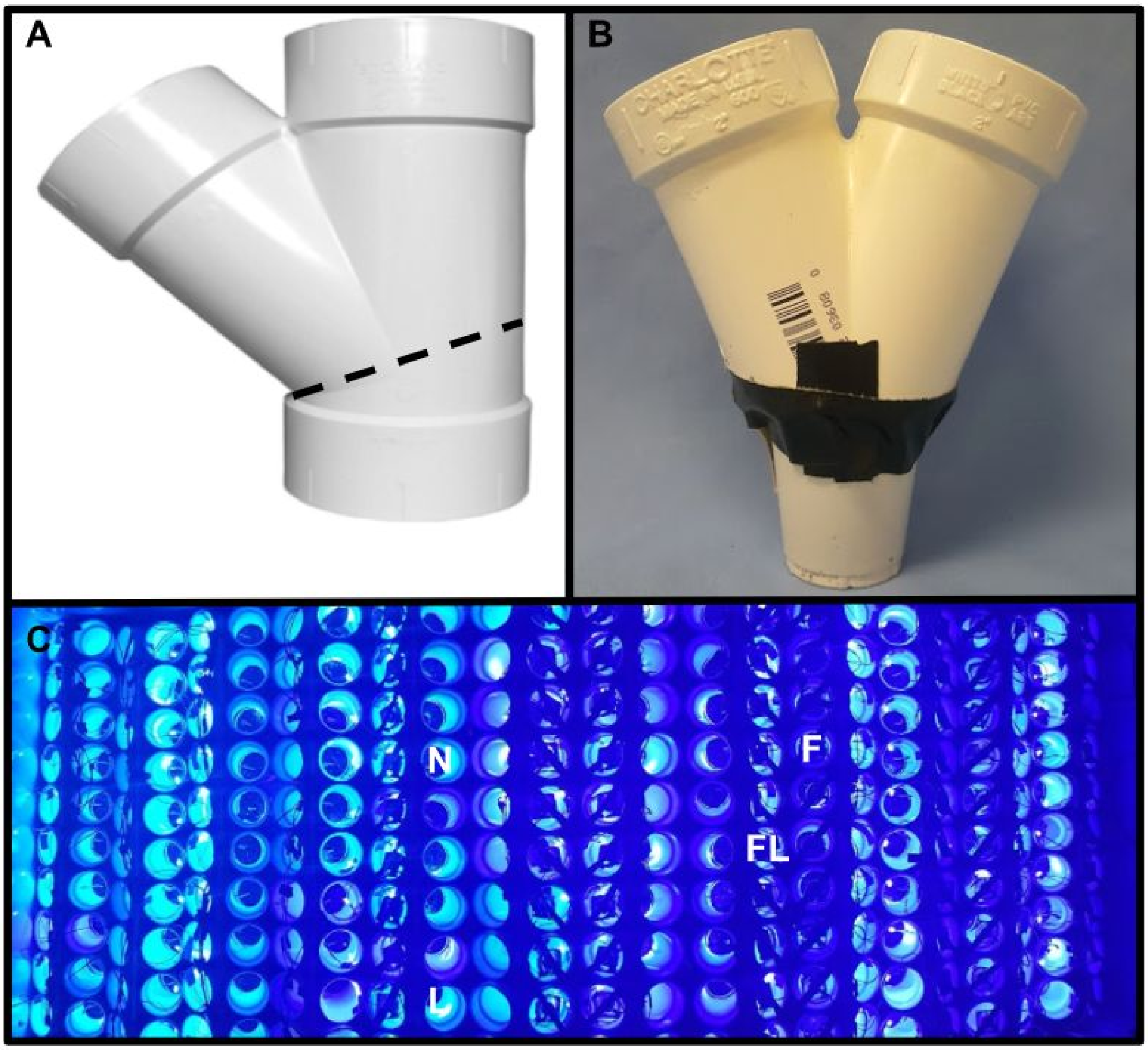


Supplemental figure 1: Y maze detail. A: Charlotte 2 in. 45 degree PVC Wye joint (Model # PVC 00600 1000) was cut at 22.5 degree angle (following the seam) using a chop saw, resulting in a bilaterally symmetrical Y shape. Maze dimensions were 55 mm base diameter, 105 mm total height, bifurcation at 65 mm height, 95 mm overall width at bifurcation point, each arm measured 60 mm diameter by 40 mm length, as per the original study. This maze is attached to the pot 1-2 days after emergence of the seedling from the soil surface. Critically, the attachment of the maze is done before condition and orientation (left vs right) is assigned to prevent bias. B: Finished Y maze attached to pot. Attachment is achieved via circumferential black tape (to prevent light leakage) with additional longer strips as needed to straighten maze as much as possible. Pot is of dimensions 5 cm diameter at top, 4 cm diameter at bottom, 6 cm height and has one central hole of approximately 2 mm diameter cut into the bottom. C: Y mazes attached to plants within controlled environment chamber. Each Y maze has two open ends, which have 4 possible electronic device configurations: N shows nothing, L shows an LED alone, F shows fan alone, and FL shows fan and LED together. These lights and fans were then moved from one maze arm to the other according to the training schedule. LEDs are attached to the lateral aspect of each Y terminus to maximize directionality of light. In these experiments, Y mazes are grouped in sets of 10 which are interconnected by a thin wooden shim which is tightly wedged between the two arms of the Y maze. This stabilizes the mazes from tipping over, a problem that plagued initial attempts.

### Scoring protocol

On testing day, during which plants must grow into one arm or the other, plants were further subdivided into control and experimental plants. All plants which had already grown into one maze arm were disqualified. Control plants were given no stimuli and left in the dark, experimental plants had the fan moved to the opposite arm from its last position and given the standard fan regimen without light. The day after testing day fans and lights were removed, plants were moved from growth chambers to a countertop and order-randomized before being scored by an independent observer with the experimenter in a separate room. Each plant was marked as either right, left, or neither. The criteria for was growth >5 mm above the decision point in only one maze arm. In the vast majority of cases, growth was in only one maze arm or neither, but two plants were disqualified and marked neither for growing into both maze arms.

### Data analysis

Once each plant’s growth direction was recorded, plant numerical IDs were matched back to their experimental condition and enantiomeric pattern of stimulus exposure (with light beginning on either the left or the right maze arm), and scored 1 if their growth direction matched the direction of most recent light exposure and 0 if it was opposite the most recent light exposure. Scores were combined for each experimental condition and the proportion of plants growing according to the phototropic expectation was established and graphed. Fisher’s exact test was used to determine different ratios between binary outcome between two conditions, and is appropriate for small samples. This is the same analysis performed by Gagliano *et al.* All tests were performed with GraphPad QuickCalcs software.

### Sample size, attrition, and exclusion criteria

Gagliano *et al.* do not mention attrition or exclusion criteria in their report, but attrition in this study was quite high at several stages, the most problematic being due to variable growth rates resulting in plants reaching the decision point prior to the test day or failing to reach it 24 hours afterwards, both of which resulted in plants being disqualified. A smaller number of plants were disqualified because their individual light or fan system was found to have disconnected during movement from one maze arm to the other, therefore interrupting their training by missing part or all of a stimulus. In these cases, plants were disqualified by the experimenter during training phases and not scored. Number of plants trained and tested vs plants successfully scored as growing left or right are shown in table 1. The controlled growth chamber used for training and testing plants had the capacity for 120 plants with mazes attached, and the experiment was performed 4 times in series to gather data. For each replication, approximately 350 seeds were germinated, of which approximately 250 would be planted, 120 transferred into the Y maze apparatus, and an average of 70.75 successfully scored as growing either left or right.

### Material differences between studies

Methods adapted from Gagliano *et al.,* with minor modifications, mostly due to differences in product availability between the United States and Australia. Gagliano *et al.* used *Pisum sativum* cv Massey Gem (source unspecified), which was not available for my lab. Green Arrow (Botanical Interests, USA) was selected as a closely related cultivar with similar growth habit. I believe this difference should not affect the results, however, because associative learning must be a complex multigenic trait which is extremely unlikely to exist within one cultivar but not be present in a very closely related cultivar. Green Arrow, like Massey Gem, is a full sun cultivar, and thus should have similar propensity for phototropism. Similarly, Gagliano *et al.* used Osmocote® seed raising and cutting mix, Scotts Australia), whereas I used Hoffman seed starter potting and planting mix (Good Earth Inc, NY, USA), though its composition closely matches the soil used in the Gagliano *et al.* experiment. Instead of one 5.3 $m^2$ controlled environment room (source unspecified), two controlled growth chambers (PGR15 Growth Chamber, Conviron) were used, one for the initial growth and soil emergence stage, one for the training and testing stage. Importantly, the light, temperature, and humidity conditions in my chambers are the same as those of Gagliano *et al.* (see Germination Conditions and Growth Conditions), and the doors to the growth chambers were only opened with the outside room darkened to prevent the interference of external light.

**Attempted pilots and troubleshooting**

The original authors did not respond to queries about the materials and troubleshooting for the experimental setup, this section is included as a summary of tips I wish I had known at the pilot stage as a resources for future experimenters. I performed two pilots during the equipment testing phase. For the first pilot, replicating the Double Light Stationary Fan condition (data not recorded because the experiment was called off) an electrical fault interrupted the training regimen. The wiring setup I used was a pair of 12V DC wires for all lights and a pair for all fans, which were each attached in parallel to their respective power wires. These power wires in turn were each connected to the output of their own 12V DC power supply, which was connected to a digital power timer programmed for the training. The solder on the 12V power supply for the lights disconnected on day 2 of the training, and the experiment was called off. The second round of pilot tested Stationary Light procedure, but at this stage the plants were in individual Y mazes not connected to each other except through the wiring system. As a result of this and the fact that the PVC mazes weigh more than the pot and soil, the apparatus was unstable, and throughout the experiment most of the plants tipped over at one point or another. They were always corrected as soon as possible, but the trauma of impact, shuffling of the soil, and gravitropism all could have a directional effect which compromised the interpretation. Any plants which spent more than several seconds on their side (those that fell when I was not present) were disqualified, but plants that fell and were immediately righted were still scored. This left 8 plants, of which 6 had grown towards the light. I accepted this as sufficient confirmation that the plants have positive phototropism, though in retrospect I wish I had replicated this condition with a larger sample size once the experimental kinks were worked out. However, the fact that plants had a statistically significant growth towards the most recent presentation of light validates positive phototropism, just not as strongly as Gagliano *et al.* reported. For the stability issue, the solution was using wooden shims to connect the plants into strips of 10 (see supplemental figure 1) and to pack the growth chamber tightly with these strips, which was sufficient to prevent any plants from tipping in the results reported in the main paper.

**Further improvements**

If the original authors stand by the work, further replication attempts are certainly in order. My hope is this work will be a source of ideas on how to further improve the study design for those that continue this work. The obvious include larger sample size and testing Massey Gem specifically, rather than a closely related cultivar. If it can be obtained, an inbred line of Massey Gem peas might reduce the variance in growth rates that caused so much attrition in this study. I'd also suggest preparing every Y maze with fan and light on each maze arm. This approach will require some more complex circuit design and double the costs for these apparatus, but would eliminate the need to move fans and lights from one side to the other during the course of the experiment. This could reduce interference with the plant growth and a substantial amount of labor. It also would make circuit failure less likely by eliminating the need to move wires during the 4 days of training and testing.
